## Supplementary material for "ARID1A governs the silencing of sex-linked transcription during male meiosis in the mouse": Suppplental Figures 1-12

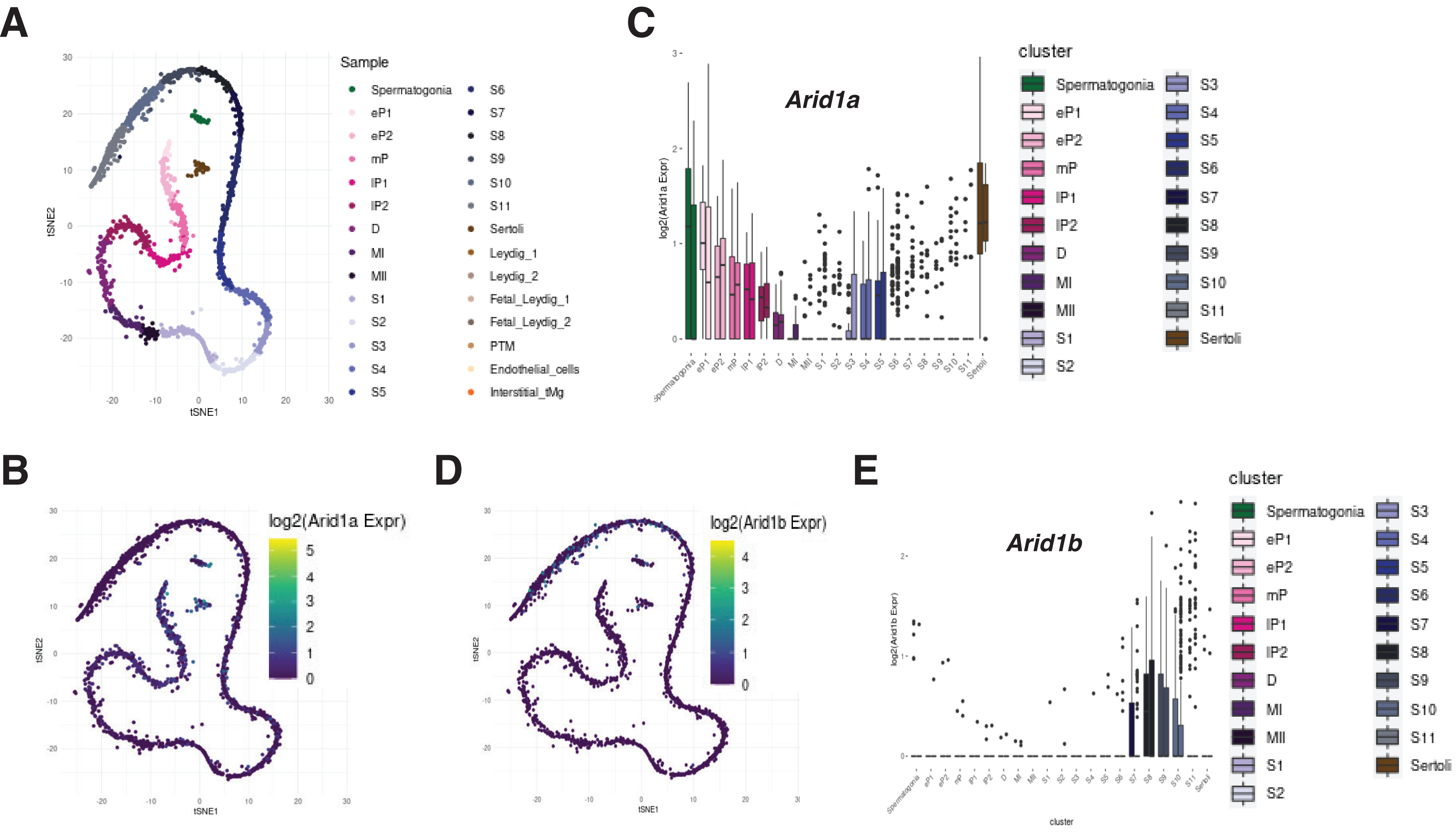
A
Sample
Spermatogonia
eP1
eP2
mP
lP1
lP2
D
MI
MII
S1
S2
S3
S4
S5
S6
S7
S8
S9
S10
S11
Sertoli
Leydig_1
Leydig_2
Fetal_Leydig_1
Fetal_Leydig_2
PTM
Endothelial_cells
Interstitial_tMg
tSNE1
tSNE2
C
Arid1a
log2(Arid1a Expr)
cluster
B
log2(Arid1a Expr)
D
log2(Arid1b Expr)
E
Arid1b
log2(Arid1b Expr)
cluster

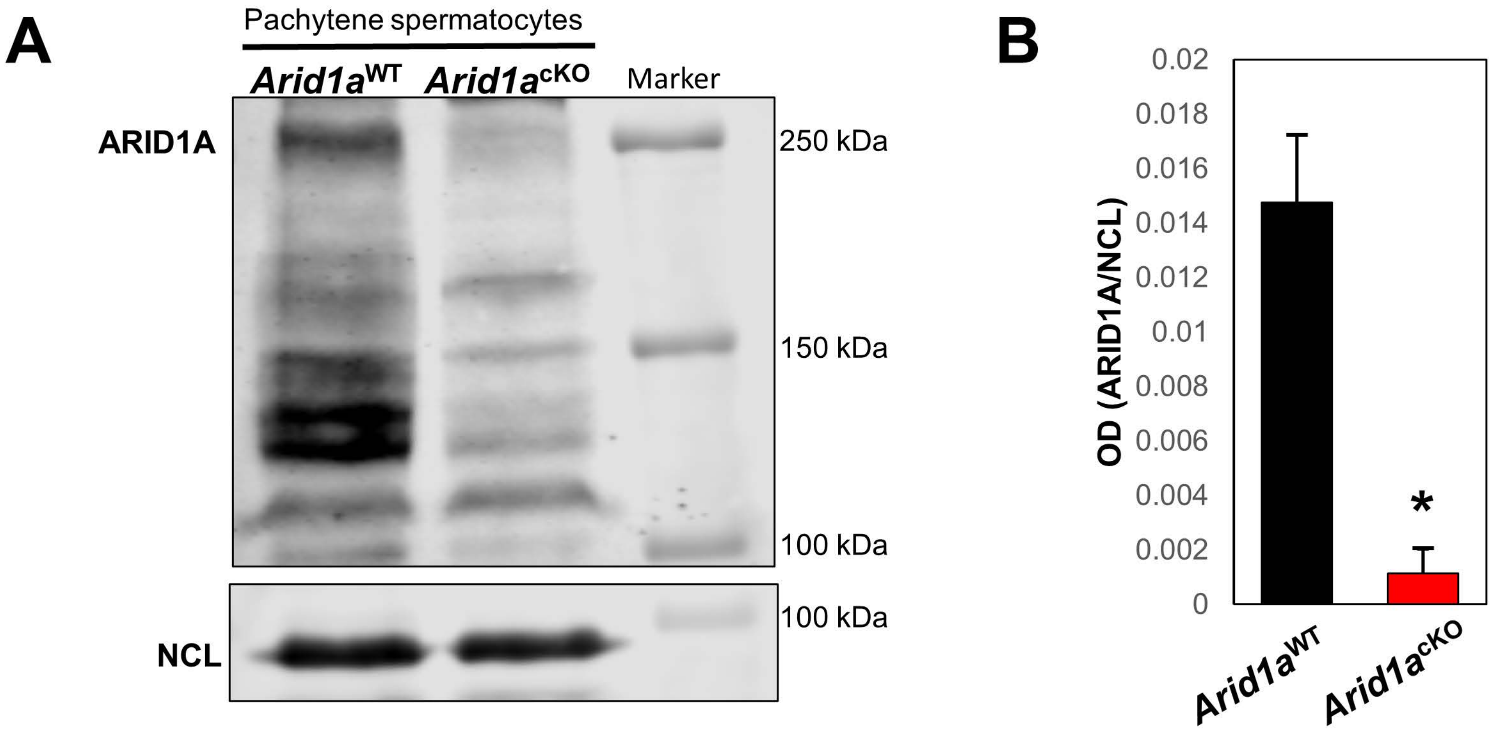
A
Pachytene spermatocytes
Arid1a^WT
Arid1a^cKO
Marker
ARID1A
250 kDa
150 kDa
100 kDa
NCL
100 kDa
B
OD (ARID1A/NCL)
0.02
0.018
0.016
0.014
0.012
0.01
0.008
0.006
0.004
0.002
0
*
Arid1a^WT
Arid1a^cKO

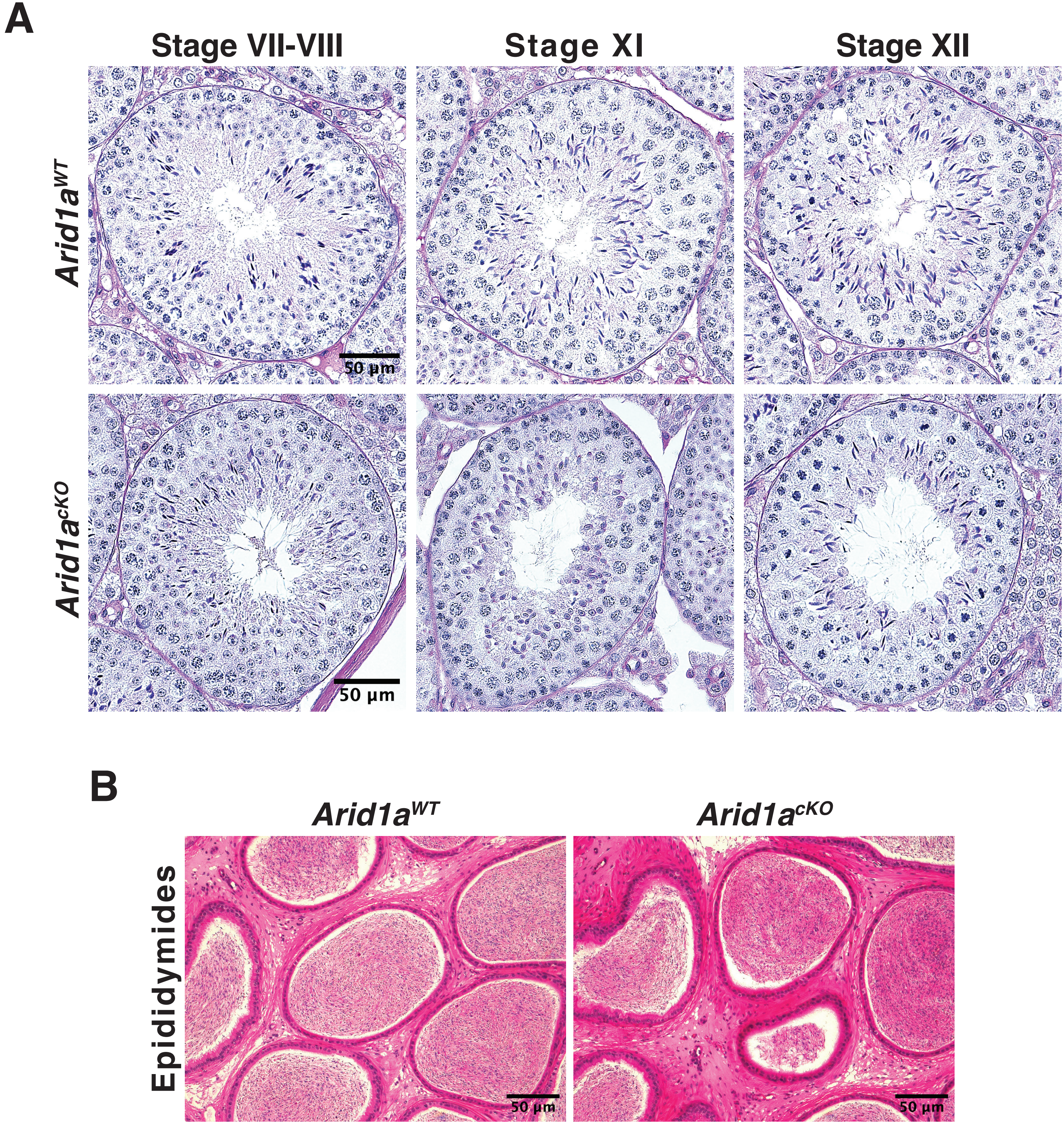

A
Stage VII-VIII
Stage XI
Stage XII
$Arid1a^{WT}$
$Arid1a^{cKO}$
50 µm
50 µm
B
$Arid1a^{WT}$
$Arid1a^{cKO}$
Epididymides
50 µm
50 µm

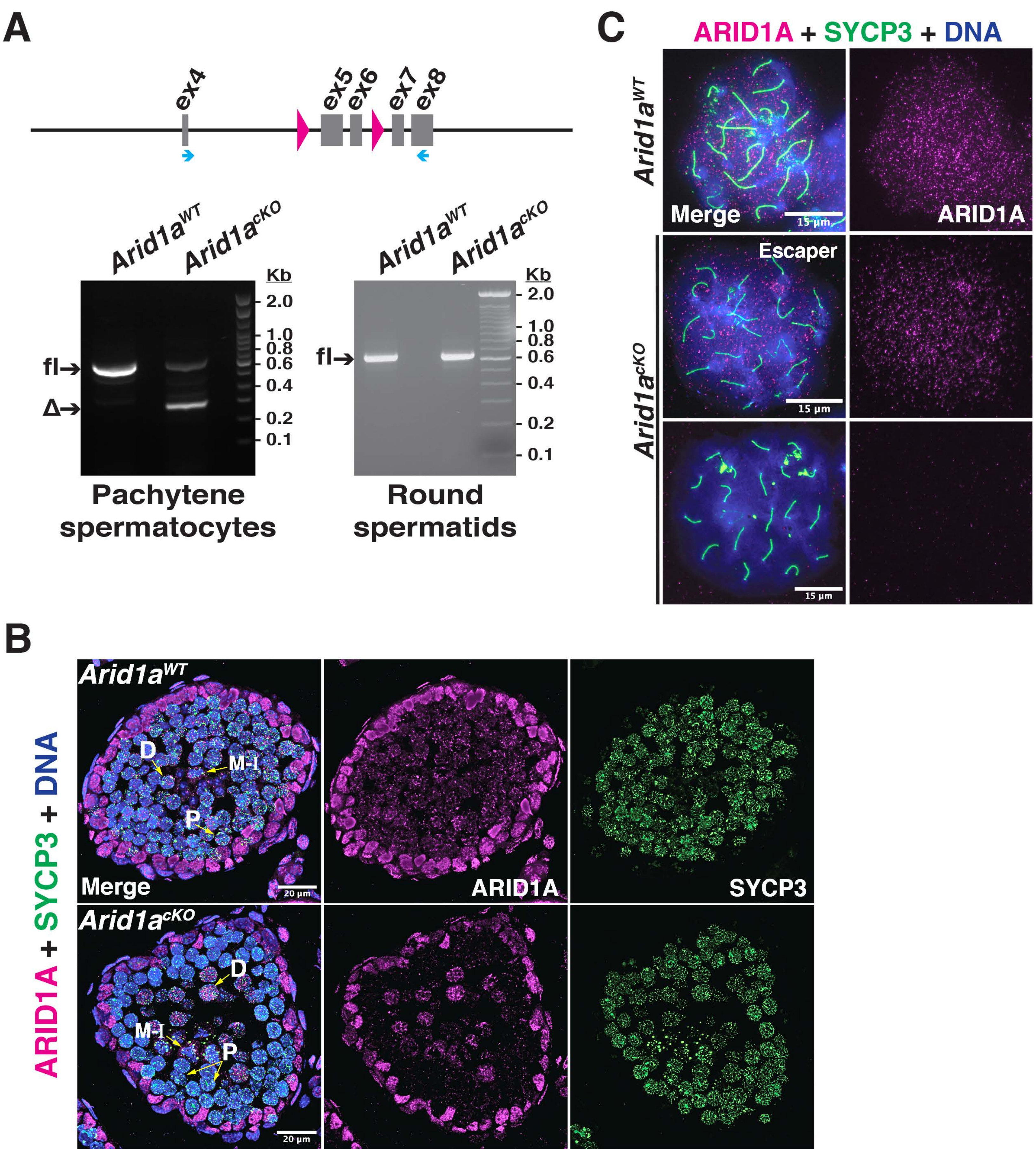
A
ex4
ex5
ex6
ex7
ex8
Arid1a^WT
Arid1a^cKO
Kb
2.0
1.0
0.8
0.6
0.4
0.2
0.1
fl→
Δ→
Pachytene spermatocytes
Round spermatids
B
ARID1A + SYCP3 + DNA
Merge
ARID1A
SYCP3
D
M-I
P
20 µm
C
Escaper
15 µm

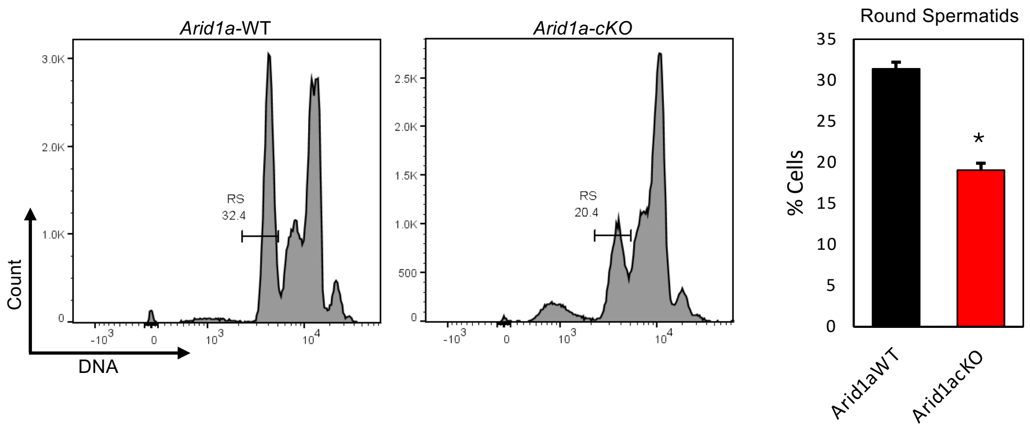

Arid1a-WT
Arid1a-cKO
Round Spermatids
RS
32.4
RS
20.4
Count
DNA
% Cells
Arid1aWT
Arid1acKO
*

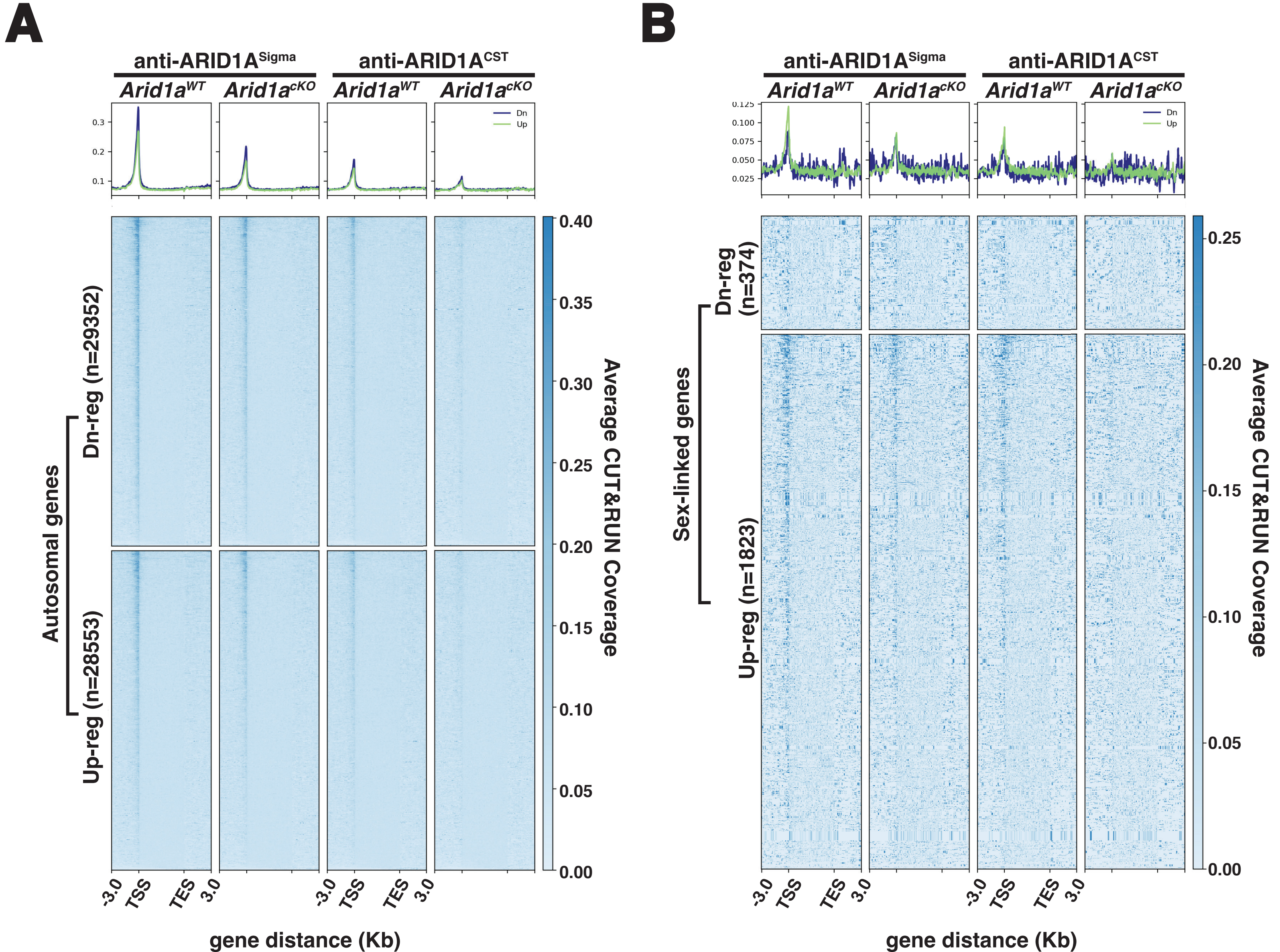
A
anti-ARID1A^Sigma
anti-ARID1A^CST
Arid1a^WT
Arid1a^cKO
Dn
Up
Autosomal genes
Dn-reg (n=29352)
Up-reg (n=28553)
-3.0
TSS
TES
3.0
gene distance (Kb)
Average CUT&RUN Coverage
B
Sex-linked genes
Dn-reg (n=374)
Up-reg (n=1823)

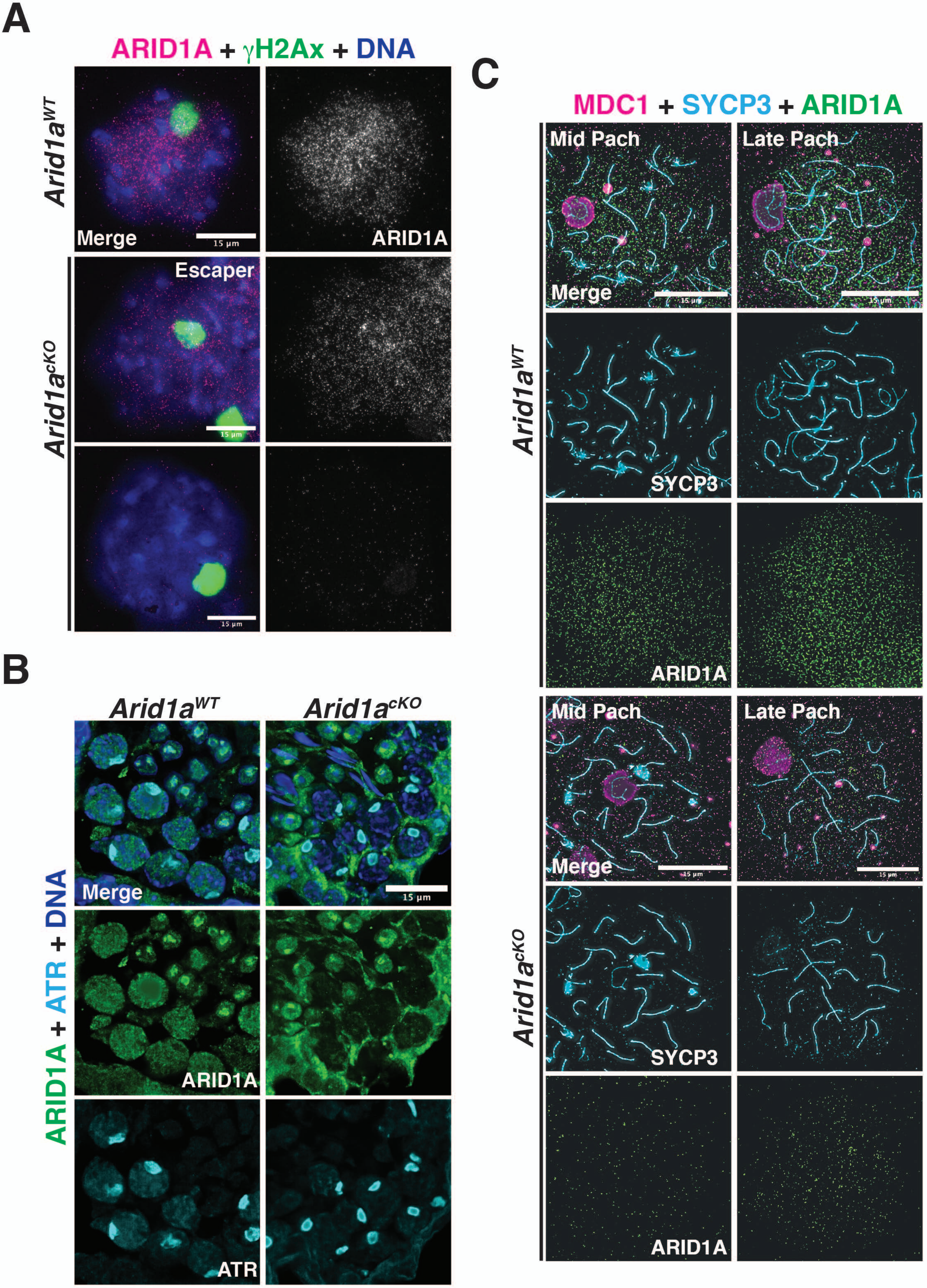

A
ARID1A + γH2Ax + DNA
Arid1a^WT
Merge
15 µm
ARID1A
Arid1a^cKO
Escaper
15 µm
15 µm
B
Arid1a^WT
Arid1a^cKO
ARID1A + ATR + DNA
Merge
15 µm
ARID1A
ATR
C
MDC1 + SYCP3 + ARID1A
Arid1a^WT
Mid Pach
Late Pach
Merge
15 µm
15 µm
SYCP3
ARID1A
Arid1a^cKO
Mid Pach
Late Pach
Merge
15 µm
15 µm
SYCP3
ARID1A

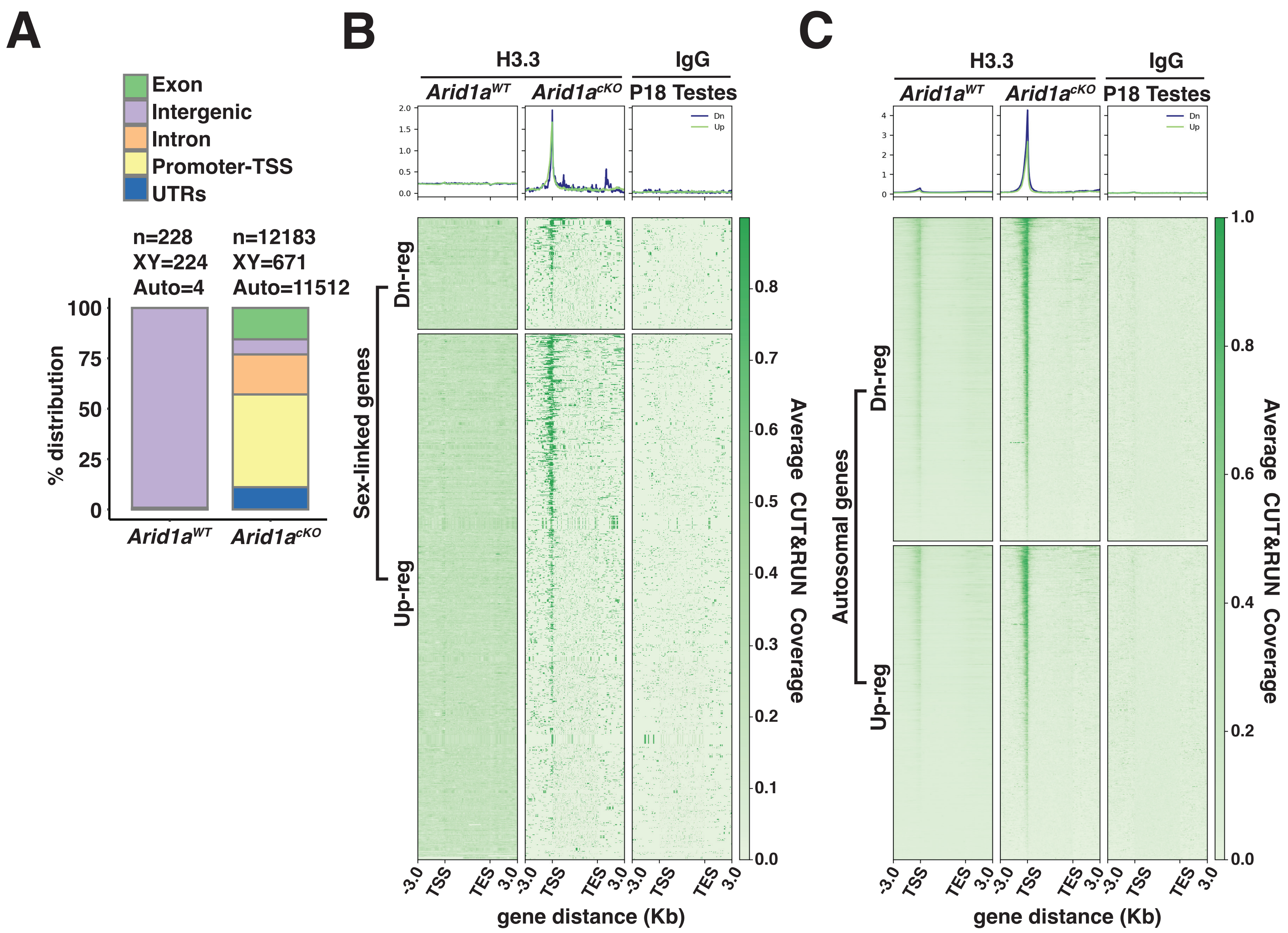

A
Exon
Intergenic
Intron
Promoter-TSS
UTRs
n=228
XY=224
Auto=4
n=12183
XY=671
Auto=11512
% distribution
100
75
50
25
0
Arid1aWT
Arid1acKO
B
H3.3
IgG
Arid1aWT
Arid1acKO
P18 Testes
Dn
Up
Sex-linked genes
Dn-reg
Up-reg
Average CUT&RUN Coverage
-3.0
TSS
TES
3.0
gene distance (Kb)
C
H3.3
IgG
Arid1aWT
Arid1acKO
P18 Testes
Autosomal genes
Dn-reg
Up-reg
Average CUT&RUN Coverage
gene distance (Kb)

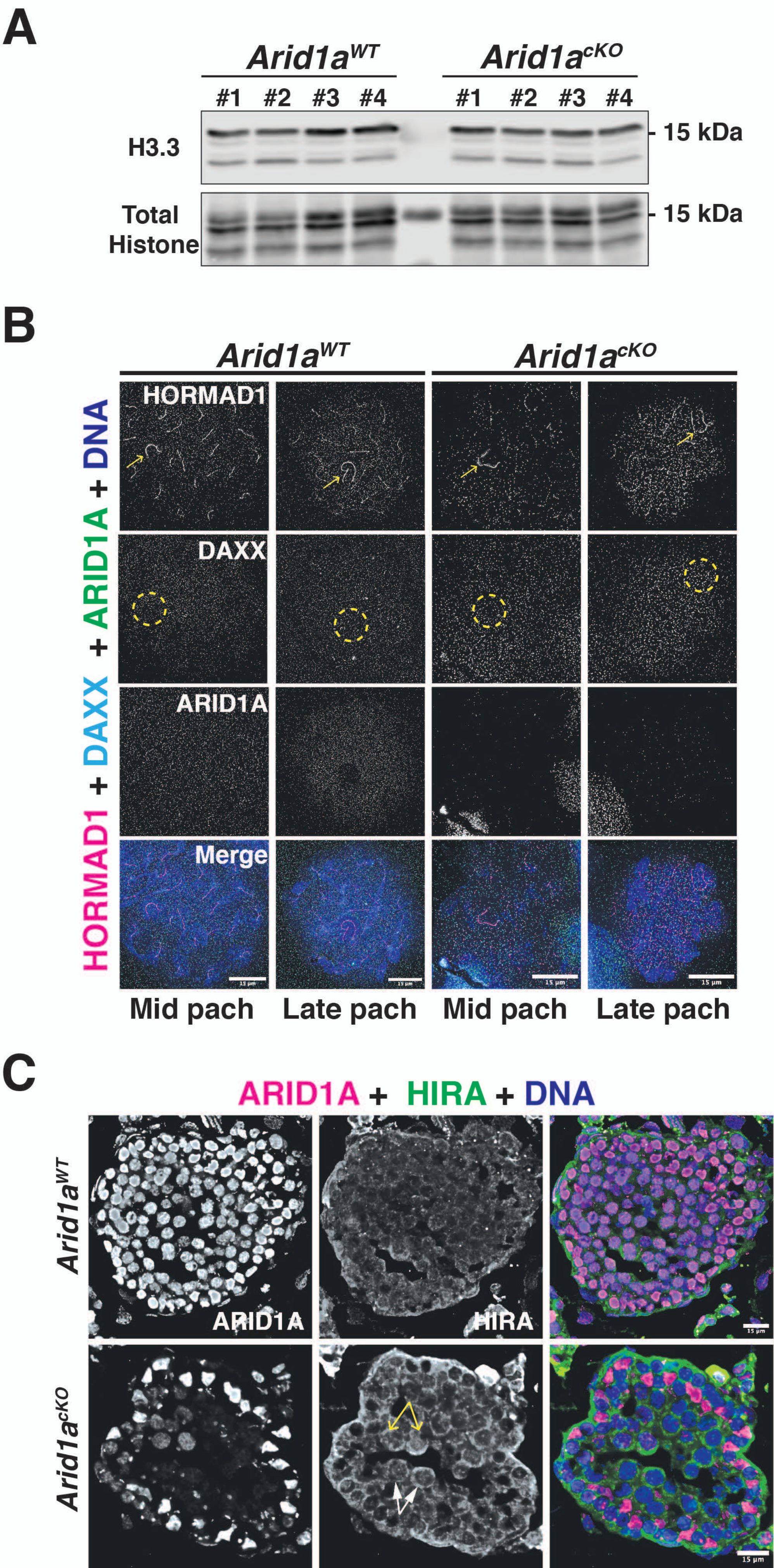
A
Arid1aWT
Arid1acKO
#1
#2
#3
#4
H3.3
15 kDa
Total Histone
B
HORMAD1
DAXX
ARID1A
Merge
HORMAD1 + DAXX + ARID1A + DNA
Mid pach
Late pach
C
ARID1A + HIRA + DNA
HIRA

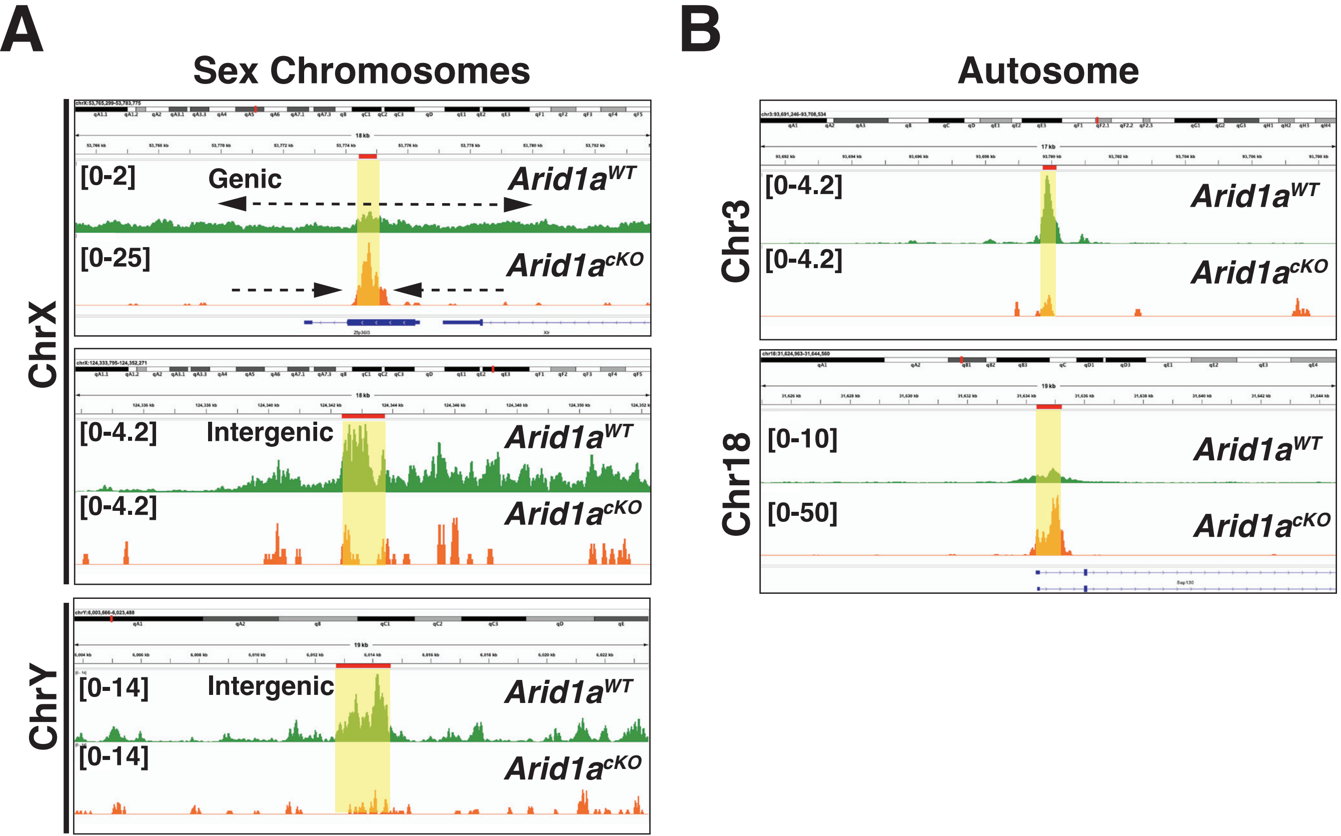

A
Sex Chromosomes
ChrX
[0-2]
Genic
Arid1a^WT
[0-25]
Arid1a^cKO
[0-4.2]
Intergenic
Arid1a^WT
[0-4.2]
Arid1a^cKO
ChrY
[0-14]
Intergenic
Arid1a^WT
[0-14]
Arid1a^cKO
B
Autosome
Chr3
[0-4.2]
Arid1a^WT
[0-4.2]
Arid1a^cKO
Chr18
[0-10]
Arid1a^WT
[0-50]
Arid1a^cKO

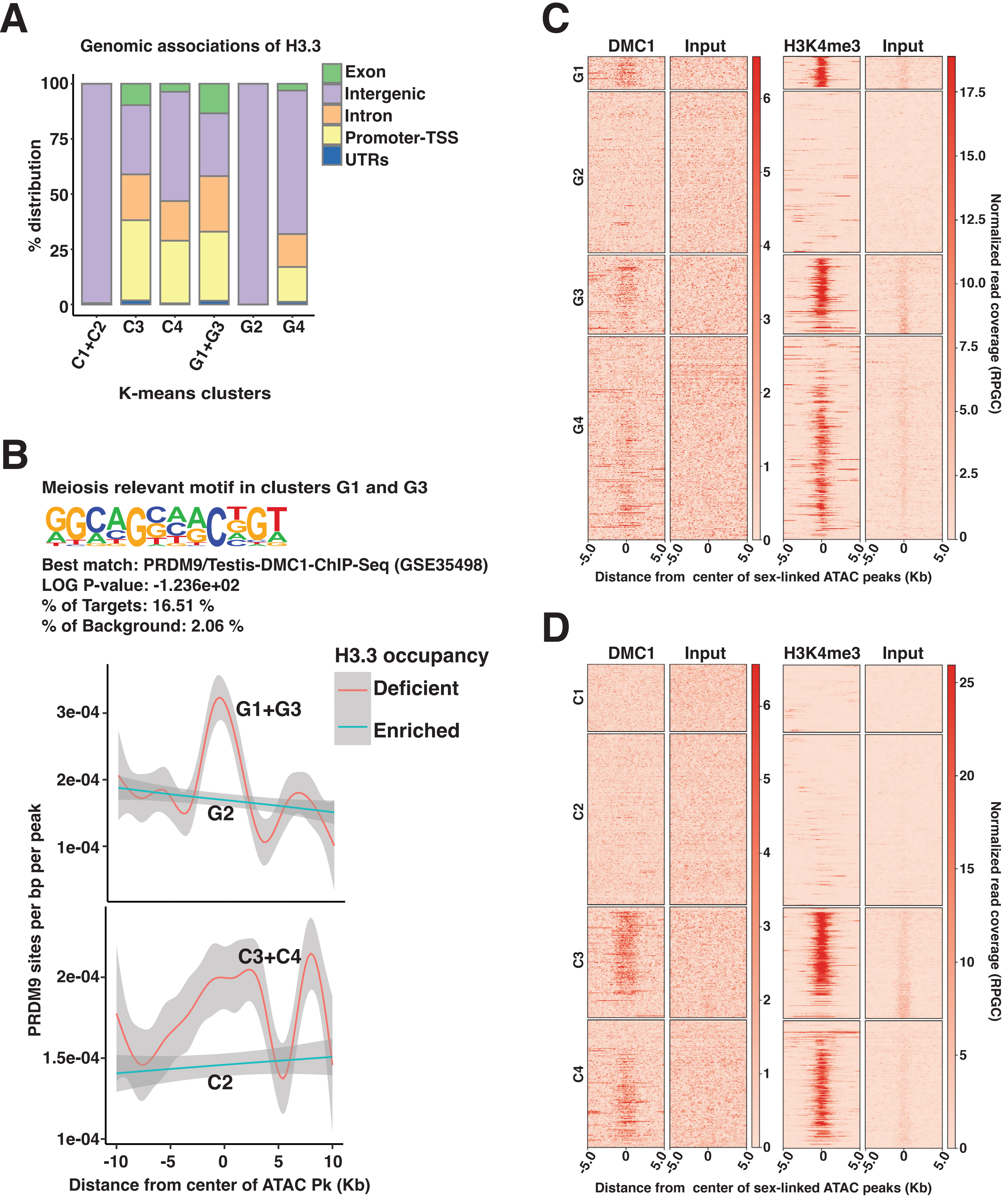
A
Genomic associations of H3.3
Exon
Intergenic
Intron
Promoter-TSS
UTRs
% distribution
100
75
50
25
0
C1+C2
C3
C4
G1+G3
G2
G4
K-means clusters
B
Meiosis relevant motif in clusters G1 and G3
Best match: PRDM9/Testis-DMC1-ChIP-Seq (GSE35498)
LOG P-value: -1.236e+02
% of Targets: 16.51 %
% of Background: 2.06 %
H3.3 occupancy
Deficient
Enriched
G1+G3
G2
C3+C4
C2
PRDM9 sites per bp per peak
3e-04
2e-04
1e-04
1.5e-04
Distance from center of ATAC Pk (Kb)
C
DMC1
Input
H3K4me3
Input
G1
G2
G3
G4
Normalized read coverage (RPGC)
Distance from center of sex-linked ATAC peaks (Kb)
D
DMC1
Input
H3K4me3
Input
C1
C2
C3
C4
Normalized read coverage (RPGC)
Distance from center of sex-linked ATAC peaks (Kb)

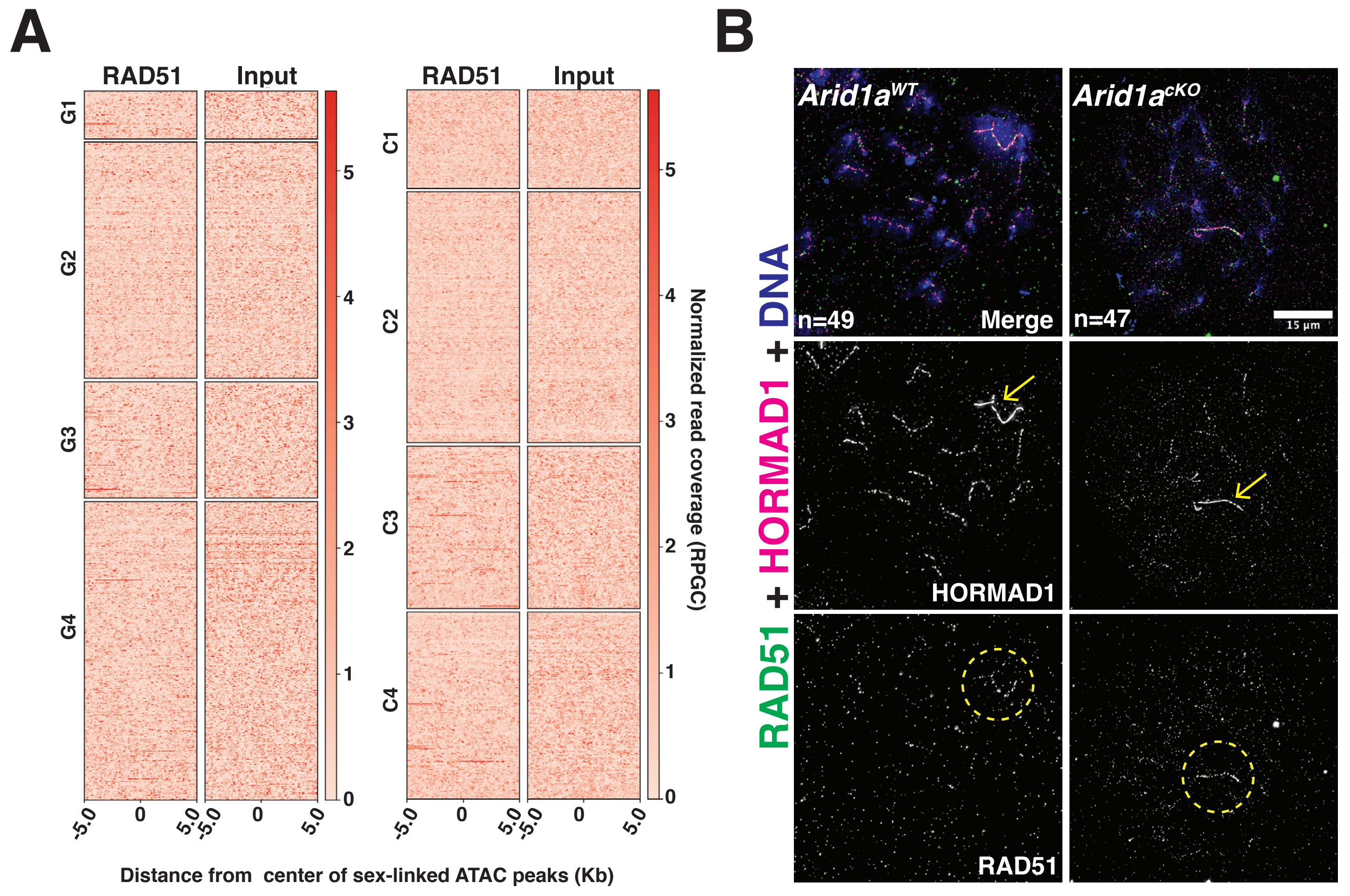
A
RAD51
Input
G1
G2
G3
G4
C1
C2
C3
C4
Normalized read coverage (RPGC)
Distance from center of sex-linked ATAC peaks (Kb)
B
Arid1a^WT
Arid1a^cKO
n=49
n=47
Merge
15 µm
HORMAD1
RAD51
RAD51 + HORMAD1 + DNA
