## Supplementary Table 1 for "ARID1A governs the silencing of sex-linked transcription during male meiosis in the mouse"

**Table S1**

| Section # | Number of PLZF+ spermatogonia | #Number of tubules examined | PLZF+  Spg/tubule | Average | SEM |
| --- | --- | --- | --- | --- | --- |
| WT1 | 648 | 91 | 7.12 | 8.87 | 1.33 |
| WT2 | 832 | 104 | 8 |  |  |
| WT3 | 782 | 68 | 11.5 |  |  |
| KO1 | 1005 | 91 | 11.04 | 10.45 | 0.49 |
| KO2 | 728 | 80 | 9.1 |  |  |
| KO3 | 883 | 82 | 10.76 |  |  |
| KO4 | 415 | 43 | 9.65 |  |  |
| KO5 | 149 | 12 | 12.41 |  |  |
| KO6 | 243 | 25 | 9.72 |  |  |

No significant difference in the numbers of undifferentiated spermatogonia expressing PLZF in 1-month-old *Arid1a^cKO^ relative to Arid1a^fl/fl^* males. SEM: Standard error of measurement.
