## Supplementary Table 3 for "ARID1A governs the silencing of sex-linked transcription during male meiosis in the mouse"

**Supplementary table 3.** List of primary and secondary antibodies

| **Antibody** | **Vendor (Catalog #)** | **Application (Dilution)** |
| --- | --- | --- |
| Rabbit anti- ARID1A | Millipore Sigma (HPA005456) | IF (1:500)  CUT&RUN (1:25) |
| Rabbit anti- ARID1A (D2A8U) | Cell signaling technology (12354) | IF (1:500)  CUT&RUN (1:25)  WB (1:1000) |
| Mouse anti- SYCP3 | Abcam (ab97672) | IF (1:500) |
| Mouse anti- γH2Ax | Millipore (05-636) | IF (1:1000) |
| Goat anti-ATR (N-19) | Santa Cruz Biotech (sc-1887) | IF (1:50) |
| Sheep anti-MDC1 | AbD Serotec (AHP799) | IF (1:500) |
| Rat anti-RNA pol II CTD phospho Ser-2 | Active Motif (61084) | IF (1:500) |
| Guinea pig anti- HORMAD1 | Gift from Dr. Atilla Töth, TU Dresden | IF (1:600) |
| Mouse anti- H3F3B (Clone 2D7-H1) | Abnova (H00003021-M01) | IF (1:200)  CUT&RUN (1:25) |
| Rabbit anti- Histone H3.3 (Clone RM190) | RevMAb Biosciences (31-1058-00) | WB (1:1000) |
| Mouse anti- Histone H3.1/3.2 | Active Motif (61630) | IF (1:500) |
| Mouse anti-DAXX | Developmental studies hybridoma bank (PCRP-DAXX-2B3) | IF (1:10) |
| Mouse anti-HIRA | Active Motif (39558) | IF (1:100) |
| Mouse anti- DMC1 | Proteintech (67176-1-Ig) | IF (1:200) |
| Rabbit anti- RAD51 | Abcam (ab176458) | IF (1:500) |
| Rabbit anti-Nucleolin | Bethyl (A300-711A) | WB (1:2000) |
| **Secondary antibodies** |  |  |
| Goat anti-mouse IgG, Alexa fluor 488 | ThermoFisher (A-11029) | IF (1:500) |
| Goat anti-rabbit IgG, Alexa fluor 568 | ThermoFisher (A-11036) | IF (1:500) |
| Donkey anti-goat IgG, Alexa fluor 633 | ThermoFisher (A-21082) | IF (1:500) |
| Donkey anti-sheep IgG, Alexa fluor 594 | ThermoFisher (A-11016) | IF (1:500) |
| Goat anti-guinea pig IgG, Alexa fluor 568 | ThermoFisher (A-11075) | IF (1:500) |
| Goat anti- rat IgG, Alexa fluor 488 | ThermoFisher (A-11006) | IF (1:500) |
| Goat anti-mouse IgG1, Alexa fluor 647 | ThermoFisher (A-21240) | IF (1:500) |
| Goat anti- rabbit IgG, Alexa fluor 488 | ThermoFisher (A-11008) | IF (1:500) |
| Goat anti-mouse IgG2b, Alexa fluor 647 | ThermoFisher (A-21242) | IF (1:500) |
| Goat anti- rabbit IgG, Alexa fluor 647 | ThermoFisher (A-21245) | IF (1:500) |
| Goat anti-mouse IgG2a, Alexa fluor 647 | ThermoFisher (A-21241) | IF (1:500) |
| IRdye 800CW Goat anti-rabbit | LI-COR (925-32211) | WB (1:10,000) |
| AffiniPure Rabbit anti-Mouse IgG (H+L) | JacksonImmunoResearch  (315-005-003) | CUT&RUN (1:100) |
| Guinea pig anti-Rabbit IgG (H+L) | Novus biologicals  (NBP1-72763) | CUT&RUN (1:100) |
